# A Reaction Norm Perspective on Reproducibility

**DOI:** 10.1101/510941

**Authors:** Bernhard Voelkl, Hanno Würbel

## Abstract

Reproducibility in biomedical research, and more specifically in preclinical animal research, has been seriously questioned. Several cases of spectacular failures to replicate findings published in the primary scientific literature have led to a perceived reproducibility crisis. Diverse threats to reproducibility have been proposed, including lack of scientific rigour, low statistical power, publication bias, analytical flexibility and fraud. An important aspect that is generally overlooked is the lack of external validity caused by rigorous standardization of both the animals and the environment. Here, we argue that a reaction norm approach to phenotypic variation, acknowledging gene-by-environment interactions, can help us seeing reproducibility of animal experiments in a new light. We illustrate how dominating environmental effects can affect inference and effect size estimates of studies and how elimination of dominant factors through standardization affects the nature of the expected phenotype variation through the reaction norms of small effect. Finally, we discuss the consequences of reaction norms of small effect for statistical analysis, specifically for random effect latent variable models and the random lab model.

## 1 Introduction

Since the mid-17th century reproducibility, i.e. the ability to reproduce an experimental outcome by an independent study is a fundamental corner stone of the scientific method which distinguishes scientific evidence from mere anecdote. In modern research, however, such independent replication has been replaced by principles of experimental design which—in principle—should render replication by independent studies redundant. In the simplest form, the effect of a predictor (independent variable) on an outcome (dependent variable) is measured in a sample of independent replicate units (individuals). Scientific evidence generated in this way is arguably reproducible if the experimental units (i.e. individuals) are true random samples of the overall target population. Despite the general wisdom, that true random samples are practically impossible to achieve when the target population is e.g. a biological species, the potential consequences of non-independence on the reproducibility of results are usually ignored. This is mirrored by the fact that no independent replication studies are generally required by funders for accepting grant proposals or by editors before accepting manuscripts for publication.

Over the last 10-15 years, however, reproducibility in biomedical research, and more specifically in preclinical animal research, has been seriously questioned [1]. Several cases of spectacular failures to replicate findings published in the primary scientific literature have led to a perceived reproducibility crisis [2, 3]. In 2011 researchers from the company Bayer reported that out of 67 in-house replication studies of published research in the areas of oncology, women’s health and cardiovascular diseases only 14 (21 percent) could fully replicate the original findings[4]. Similarly researchers of the company Amgen have replicated 53 original research studies deemed ‘landmark’ studies in haemathology or oncology, recovering the original findings only in 6 cases (11 percent)[5]. These reports and a surge of meta-analyses confirming low replication rates (e.g. [6–8]) lead to a heated debate within as well as outside the scientific community about the usefulness of animal models for bio-medical research [3, 2, 9–11].

Several potential causes for poor reproducibility have been proposed, including lack of scientific rigour, low statistical power, publication bias, analytical flexibility, and perverse incentives in research—leading in some cases to outright fraud [10, 2, 3]. While all of these aspects might contribute to replication failure, we will here focus on another aspect that is all too often ignored: biological variation. Biological variation is the sum of genetic variation, environmentally induced variation and variation due to the interaction between environment and genotype (G × E interaction). As the response of an animal to an experimental treatment (e.g. a drug) depends on the phenotypic state of the animal, the response, too, is a product of the genotype and the environmental conditions. Despite attempts to standardize animal facilities, laboratories always differ in many environmental factors that affect the animals’ phenotype (e.g. noise, odours, microbiota, or personnel [12–15, 13, 16]). In a landmark study Crabbe and colleagues [12] investigated the confounding effects of the laboratory environment and G × E interactions on behavioural strain differences in mice. Despite rigorous standardization of housing conditions and study protocols across three laboratories, systematic differences were found between laboratories, as well as significant interactions between genotype and laboratory. Even temporal variation within a single laboratory can lead to relevant effects, as demonstrated in a recent study where researchers found considerable phenotypic variation between different batches of knockout mice tested successively in the same laboratory [17, 18].

The reaction norm is a concept helping to explain the observation that individuals of the same genotype will produce different phenotypes if they experience different environmental conditions [19]. It is the result of a complex environmental cue response system, which buffers the functioning of the organism against environmental and genetic perturbations [20–22]. The consequence of such a regulatory system is that environmental influences can play an important part in shaping the phenotype. Environmental influences do not only play a role at the time of assessment of the phenotype but throughout the ontogeny of the organism [23]. A reaction norm perspective on phenotypic traits unifies two concepts which have often been treated as opposing mechanisms: phenotype diversification due to environmental variation (plasticity) and the limitation of phenotypic variation by mechanisms that buffer development against genetic and environmental variation (canalization). Both plasticity and canalization have been considered as adaptive traits evolved as a consequence of environmental variation, though following Woltereck’s [19] arguments, it is the reaction norm itself that one should consider as the evolved trait [24]. Its adaptive value is, however, limited to a certain range of environmental variation: environmental situations that lie far outside the range of environments a species experienced over its evolutionary past, can overtax the organism’s ability to appropriately respond to the situation and lead to maladaptive or pathological responses. With respect to reproducibility it must be emphasised that ‘phenotype’ is not restricted to visible differences between individuals but does equally refer to differences in physiological or behavioural responses to any sort of stimulation or treatment.

We have recently argued that a failure to recognize the implications of reaction norms might seriously compromise reproducibility in bioscience—specifically in in-vivo research [25, 26]. Laboratory experiments that are conducted with inbred animals under highly standardized conditions are testing only a very narrow range of one specific reaction norm. Independent replicate studies that fail to reproduce the original findings might not necessarily indicate that the original study was poorly done or reported, but rather that the replicate study was probing a different region of the norm of reaction [27]. Therefore, the attempt to improve reproducibility through rigorous standardization of both genotype and environment has been referred to as “standardization fallacy” [28]. Here we will explore this proposition in more detail, first consider the case of a single dominating environmental factor, and then the reaction norms of small effect. In practical terms this will lead us to emphasise the importance of including the laboratory environment as a factor in multi-laboratory studies and meta-analyses or to consider introducing a correction factor in the statistical model to account for predicted between-lab variation.

## 2 Conceptualizing The Reaction Norm

The reaction norm can be conceptualized as a function mapping an environmental parameter to an expected value of a phenotypic trait (Figure 1).

**Fig. 1.**
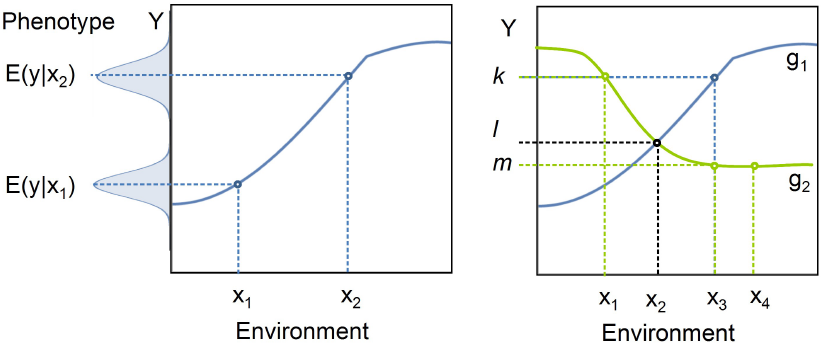
(a) The reaction norm allows describing the relationship between the expected value of a phenotypic trait (*E*(*Y*)) and an environmental parameter (*X*) for a specific genotype. The observed values of the phenotypic state (indicated by the Gaussian bell curves) will vary due test variation, measurement error, and due to biological variation induced by variation in other environmental parameters. (b) The reaction is a genotype specific property: different genotypes (*g*_1_,*g*_2_) can have different reaction norms, with the effect that for the same environmental parameter value, *x*_3_, *g*_1_ and *g*_2_ produce different expected trait values, *k* and *m*. For some *x*, both genotypes can have the same expected values for *y* (e.g. *E*(*y|g*_1_*, x*_2_) = *E*(*y|g*_2_*, x*_2_) = *l*) and different genotypes can have the same expected trait value under different environmental conditions (e.g. *E*(*y g*_2_*, x*_1_) = *E*(*y g*_1_*, x*_3_) = *k*). If the reaction norm is flat, we expect the same trait value even under different environmental conditions (e.g. *E*(*y|g*_2_*, x*_3_) = *E*(*y|g*_2_*, x*_4_) = *m*).

If we denote the environmental parameter as *X* and the phenotypic trait of the organism as *Y*, then the norm of reaction *h*(·) gives the expected value for *Y* given the environmental state *x* as *E*(*y|x*) = *h*(*x*). In many cases the phenotypic trait will be a continuous valued trait. In this case we can describe the distribution of expected values for the trait by a probability density function (PDF) *f* (*y*). The environmental parameter is assumed to be a characteristic that can be measured on a continuous scale. Environments differ in the environmental parameter and the probability of finding the environment in a specific state regarding this parameter can be given by a probability density function *g*(*x*). Hence, with the help of the reaction norm, we can describe the relationship between the expected trait value and the distribution of the environmental states with the composite function

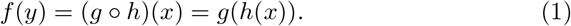

Originally Woltereck [19] referred to the relationship between a specific environmental variable and the phenotype as *Phänotypenkurve* (phenotype curve), while he used the term *Reaktionsnorm* (reaction norm) for specifying the collective influence of all environmental variables. However, later Woltereck widened the use of the term reaction norm to include also small subsets of phenotype curves or even phenotype curves of a single environmental variable. Today the term norm of reaction is usually used to describe the relationship between a single environmental parameter on the expected phenotype of the organism [29, 30]. In evolutionary ecology reaction norms are often the target of the study. Reaction norms are studied experimentally by systematically varying one environmental parameter. If one wants to describe the combined effect of two or more environmental parameters on the phenotype, the norm of reaction takes on the form of a surface or a hypersurface. Conceptually, there is no bound for the number of dimensions included, though limits of human imagination sets constrains as the heuristic value of the model quickly decreases with increasing dimensionality. Furthermore, collecting empirical data becomes very cumbersome when combinations of several parameters need to be varied systematically. For these two reasons defining high-dimensional norms of reaction is an approach rarely taken or advised.

### 2.1 Dominating Factors

In most cases of biomedical research, environmentally induced trait variation appart from the treatment effect is not of interest and considered as unwanted noise. The predominant approach taken to deal with environmentally induced variation is to identify potential dominating environmental parameters and keep them constant (standardization). Where, we speak of a parameter as ‘dominant’ if it contribute much more to the total environmentally induced trait variance than most other parameters. In those cases, where a dominating factor can be identified but not controlled, it might be recorded and added to the analysis as co-variate or nuisance factor (Figure 2). The very idea of environmental standardization is, thus, to reduce environmentally induced trait variation by reducing variation of all those environmental factors that are known to—or are suspected to—cause trait variation. The list of factors standardized in most pre-clinical studies with rodent model organisms includes (but is not limited to) cage size, cage content (nesting material, shelter, enrichment devices), housing temperature, humidity, light regime, stocking density, food and water supply, handling techniques and cage maintenance routines. In fact, even many more environmental factors are standardized, though some of them seem to be so self-evident or trivial that they are hardly ever mentioned and easily overlooked (e.g. all laboratory environments are free of catastrophic events like hailstorms or feline predators). Thus, rigorous standardization is presumed to eliminate most or all dominating factors and, hence, lead to a substantial reduction of environmental variation and arguably also to a reduction in environmentally induced trait variation. Study-specific standardization will mainly reduce within-study trait variation, while standardization across studies (harmonization) will reduce both within- and between-study variation.

**Fig. 2.**
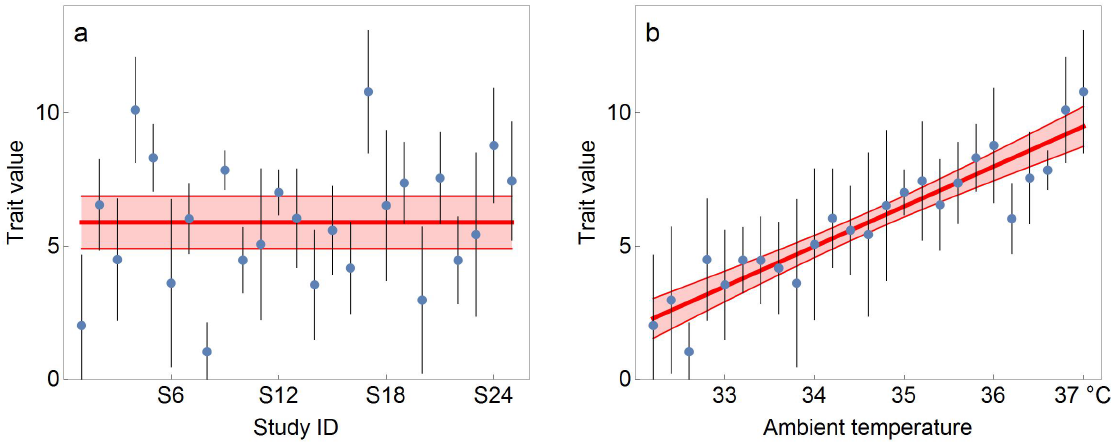
The effect of dominating factors on effect size estimates and reproducibility. Panel (a) shows the hypothetical results of 25 studies, where between-study variability is relatively large in comparison to within study variability and the confidence intervals of several studies would not include the summary effect size estimate. In panel (b), however, studies are sorted by an environmental gradient (ambient temperature) on the y-axis, suggesting that this environmental factor has a linear influence on the effect size of the experimental treatment. In this case, inclusion of this factor, would allow giving predicted values with respect to the environmental variable and most studies capture the predicted value for the respective ambient temperature. In the case of a specific environmental factor that was reliably measured and reported for all studies, such a regression approach would, indeed, be the best option for both estimating the conditional effect size and estimating replication success.

### 2.2 Reaction Norms of Small Effect

If all environmental factors with dominating contributions to trait variation have been “neutralized” in a big sweep, one might believe that the remaining environmentally induced variation is of little interest. This, however, might not necessarily be the case, because in addition to environmental conditions, the genetic background of the laboratory animals is also highly standardized when experiments are conducted with inbred mouse strains. Mice used in a single study will be delivered from the same breeding facility and stem from the same breeding line. As a consequence, individual genetic variation is very small, with the result that environmentally induced variation and G E interactions might still make up most of the total biological variation of the organism [28]. Environmental effects should, therefore, still be taken into account. Yet, the nature of the combined environmental influences has changed. Originally, we were confronted with the situation of many environmental parameters having a small effect on trait variation and one or a small number of dominating parameters, contributing much more to trait variability. However, after dominating factors have been taken care off, we should be left only with a large number of factors, each having a small effect on the total variance. This situation requires a different treatment. Assuming that those factors are additive and independent of each other and recalling the central limit theorem [31–33], we can expect that under those assumptions the limiting distribution for the effect of the environmental states can be described by a Gaussian random variable 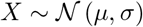.

## 3 Reproducibility

As the reaction norm allows relating environmental variation to expected variation of trait values *Y*, we might ask, whether this can help us in defining an acceptance region, in which the effect size estimate of a replicate study has to fall, in order to be considered a ‘successful’ replication. Traditionally, the discussion how to find this region has focused almost exclusively on the domain of *Y*—the trait—by partitioning the observed variation in the trait value in variance attributed to laboratory (i.e. environmental variation) and variance attributed to individual variation and measurement error. Here, we suggest a conceptually different approach: instead of defining the acceptance region based on observed trait variation, we want to define the acceptance region based on the range of expected values given the environmental states. We can consider two different scenarios: (*a*) the reaction norm is known and the value *x* for the environmental variable for the specific studies are known, and (*b*) the reaction norm is known and the distribution for the environmental variable is known. Under scenario *a* we can use the reaction norm to find the expected value for *y* for the original study as *E*(*y|x*_1_) = *h*(*x*_1_), where *x*_1_ is the value for the environmental variable of the original study. Likewise, the expected value for *y* of a replicate study done under environmental condition *x*_2_ is given by *E*(*y|x*_2_) = *h*(*x*_2_). Different measures for reproducibility have been suggested, though for our purpose a very simple definition might suffice. We say that a replication study successfully reproduced the original finding if its parameter estimate falls within the confidence interval of the original study. The replicate study can be said to reproduce the findings of the original study if

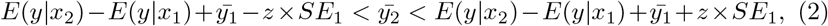

where 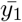 is the mean of the observed values of the original study, 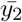 is the mean of the observed values of the replicate study, *SE*_1_ is the standard error for the mean estimate of the original study done under environmental condition *x*_1_, and *z* is a parameter determining the confidence level. In words, as we know the difference in expected trait values for the environmental conditions under which original and replicate study have been conducted, we can shift the confidence interval for the mean estimate of the original study by that amount before testing whether the mean estimate of the replicate study falls within that interval. Practically, such cases where the reaction norm and the environmental parameters are known might be rare, because if they are known, then the expected value for *y* could be deduced readily from *x* and there would be little need to actually perform the experiment. Under scenario *b* the reaction norm is known but the researcher is blind to the actual parameter values for the environmental variable *X* under which the original study or the replication study were performed. However, the researcher knows the overall distribution for *X*. In this case we can approach the question of reproducibility differently. If we know the distribution for *X* and the reaction norm *h*(·), equ. 1 allows us to evaluate the distribution for the expected values for *Y*. We can use this distribution to ask for the likelihood that the mean value from an observed set of values, 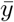, could stem from *Y* by calculating the probability that a randomly drawn value from *Y* would be more extreme than 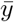. We can do this for both, the observed mean for the original study 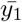 and the observed mean for the replicate study 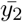. If the product of those probabilities is sufficiently large (lager than a critical value *L*), we have no reason to reject the idea that both estimates faithfully reflect randomly sampled realizations of the environmental parameter *X*. For 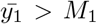 and 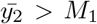, where *M*_1_ is the first moment of *f* (·), we can speak of successful replication if

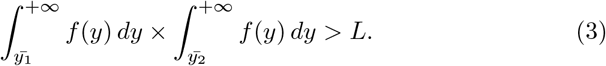

In case of 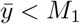 the respective integral is to be taken from 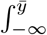. Like scenario *a*, scenario *b* suffers from the problem that the reaction norm must be known. If it is not known, we cannot proceed this way, but the reaction norms of small effect can at least be integrated in the statistical model. For this case we noted that the combined effect of many environmental variables should result in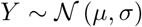. The contribution of the reaction norms of small effect to an observed difference between two study outcomes will be confounded with other sources of between-study variation, thus we cannot isolate it and consequently also not determine its effect on reproducibility. However, the reaction norms of small effect can be subsumed in the random variable for lab or study in a latent variable model and, hence, statistically taken care of.

## 4 Random Lab Model

A statistical approach incorporating the reaction norm into estimates of individual studies has been suggested by Kafkafi and colleagues [34], dubbed the random lab model (RLM). This model adds ‘noise’ for the presumed variation contributed by the *G × E* interaction term to the individual variation, generating an ‘adjusted yardstick’ for inference and parameter estimates. It is, thus, raising the benchmark for finding significant results by trading statistical power for increased realism through wider confidence intervals of the effect size estimates. The effect of this adjustment is technically achieved by adding a penalizing *G × E* term to the variance. The standard error for the effect size estimate of a simple contrast of two groups (e.g. ‘test’ and ‘control’) can, then, be calculated as

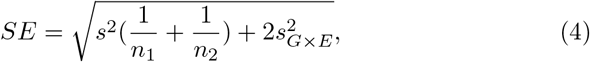

where *s*^2^ is the observed variance, *n*_1_ and *n*_2_ are the respective sample sizes for treatment and control groups, and 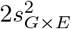 is the added *G* × *E* noise’ [34]. The latter term cannot be estimated from data from a single experiment, but it is suggested—or hoped for—that large data bases or meta analyses will allow giving rough approximate values for specific fields of research and specific types of interventions.

## 5 Discussion

We started off with the observation that the phenotype of an organism is always a product of its genotype and the environmental circumstances under which it developed. Thus, a phenotypic trait should not be considered as a fixed entity but as a conditional property of the organism. Experimenters have long identified environmental clustering—be it as sites, laboratories, batches, racks, cages—as potential sources for covariation. The seemingly logical solution to this problem is, to add shared environment as a random effect in the statistical model. For example, if a large biomedical intervention study is carried out at several laboratories, then a joint analysis would include the identity of the laboratory as a random factor in the analysis. In single-lab studies batch or cage are often added as random factors. These random factors are by default modelled as normally distributed random variables. As several authors have noted (e.g. [35–38]) this assumption might often be made for computational convenience and not because of compelling empirical evidence. From a conceptual viewpoint it is not always justified: it might work well if the environmental influence is a sum of many different underlying processes (reaction norms of small effect), while the presence of dominating factors can lead to non-normal distributions for the expected trait value.

Next, we have noted that reaction norms come in two flavours: dominating factors and factors of small effect. Given the usually continuous nature of environmental effects on trait values, this is a rather arbitrary distinction that would defy any attempt of operationalization. Dominating factors are environmental factors that contribute much more to the overall trait variation, than other environmental factors, but for practical purposes we can simply define dominating factors as factors where we can see clear effects on the trait variation given realistic (reasonably small) sample sizes. If such effects exist, vigilant experimenters will either control the environmental parameter (keeping it constant) or incorporate it in the analysis by systematically varying it and adding it to the model. Furthermore, we can expect a large number of environmental parameters having a small effect on the expected phenotype value. Employing the central limit theorem, we suggested summarizing the effects of all those parameters in a single normally distributed random variable. The question arises, whether those small environmental effects can have an effect on the reproducibility of a study result. We argue that this can, indeed, be the case for two reasons. First, even if the effect of a single environmental parameter might be rather small, the combined effects of many such parameters can—sometimes—become substantial. (Though in many cases it will not as a result of the regression to the mean.) Second, what we see in biomedical research is a tendency for standardizing many aspects of experimental studies. Standardizing instruments and measurement protocols means reducing measurement error. Standardizing housing conditions and testing conditions means eliminating most dominating environmental factors and, hence, reducing the overall variation. At the same time, standardizing the genotype by working with highly inbred lines means that also the genetic variation is largely reduced—leading again to a reduction of variance of the phenotype. Thus, while the overall phenotypic variation is reduced through standardization, the relative proportion of the phenotypic variation contributed by the remaining environmental factors will consequently increase [28]. As the reduction of measurement error and genetic variation results in a larger proportion of phenotype variation that can be attributed to the reaction norms of small effect, we have to consider what consequences this has for the distribution of the expected trait value.

From viewing between-study variation from a reaction norm perspective, we can learn two important things. First, as soon as the slope for the reaction norm is not flat, the environment affects the expected trait value and should be incorporated in any explanatory model as latent variable. In analyses of multi-lab studies and in meta-analyses this is done by treating the laboratory, the study site, or the study as random factor of a mixed effect model. Indeed, over the last decades several authors have emphasised and diligently advocated the use of mixed effect models for multi-centre studies [39, 40] and meta analyses [41]. Their efforts have not been in vain and today mixed effect models can be considered the standard approach to dealing with lab-to-lab or clinic-to-clinic variation. However, while those recommendations for the use of mixed effect models were based on statistical arguments (non-independence and the observation that adding a random factor for lab or clinic can reduce the unexplained error term), we arrived at the same suggestion from—what we would call—first principles of biology: the norm of reaction as a cogent product of stabilizing selection. Second, as soon as dominating factors have non-linear reaction norms, it becomes likely that the resulting distribution for expected trait values is not normal. Does this mean that multi-centre studies or meta-analyses implicitly assuming a normally distributed latent variable for the combined effects of lab environment are wrong? From a conceptual viewpoint, this might indeed be a questionable assumption; however this might not matter too much for practical purposes. For most statistical models it is sufficient that normality is only approximately met as the algorithms might be rather robust against moderate deviations from normality [38, 42–44]. That is, if the reaction norm for the dominating factor does not lead to a heavily skewed or distorted distribution of the latent variable, then the effect on the model outcome might be negligible. If one has reason to believe that the assumption is substantially violated, then a non-parametric modelling approach based on mixture-models [36, 35] or Markov chain Monte Carlo methods [45] might offer suitable alternatives.

## 6 Conclusion

When studying living organisms we are faced with inherent biological variation which is distinct from random noise or measurement error and which is fundamental to the correct interpretation of experimental results. Fully acknowledging this requires adopting a reaction norm perspective on physiological and behavioural responses. This will lead to a re-thinking of parameter estimation and inference, it will let us see reproducibility in a new light and it can even help gaining new insights into adaptive responses and gene-by-environment interactions. Here, we have tried to dissect its implications for the reproducibility debate and, more generally, what it means for the interpretation of experimental results in biomedical research.

## Acknowledgements

Funding through the Swiss National Science Foundation, SNF grant no. 310030-179254 to HW. We thank four anonymous reviewers for their helpful comments on an earlier version of this manuscript.

## Notes

### Competing Interest Statement

The authors have declared no competing interest.

### Summary of Updates

Substantial revision

